# The Laplacian eigenmaps dimensionality reduction of fMRI data for discovering stimulus-induced changes in the resting-state brain activity

**DOI:** 10.1101/2020.12.10.419861

**Authors:** Nikita Pospelov, Alina Tetereva, Olga Martynova, Konstantin Anokhin

**Affiliations:** Institute for Advanced Brain Studies, Lomonosov Moscow State University, Moscow, Russia; Institute of Higher Nervous Activity and Neurophysiology of the Russian Academy of Science; Department of Psychology, University of Otago, Dunedin, New Zealand

**Keywords:** Resting-state fMRI, BOLD signal, conditioned fear, dimensionality reduction, Laplacian eigenmaps, multiresolution analysis

## Abstract

The resting brain at wakefulness is active even in the absence of goal-directed behavior or salient stimuli. However, patterns of this resting-state (RS) activity can undergo alterations following exposure to meaningful stimuli. This study aimed to develop an unbiased method to detect such changes in the RS activity after exposure to emotionally meaningful stimuli. For this purpose, we used functional magnetic resonance imaging (fMRI) of RS brain activity before and after the acquisition and extinction of experimental conditioned fear. A group of healthy volunteers participated in three fMRI sessions: a RS before fear conditioning, a fear extinction session, and a RS immediately after fear extinction. The fear-conditioning paradigm consisted of three neutral visual stimuli paired with a partial reinforcement by a mild electric current. We used both linear and non-linear dimensionality reduction approaches to distinguish between the initial RS and the RS after stimuli exposure. The principal component analysis (PCA) as a linear dimensionality reduction method did not differentiate these states. Using the non-linear Laplacian eigenmaps manifold learning method, we were able to show significant differences between the two RSs at the level of individual participants. This detection was further improved by smoothing the BOLD signal with the wavelet multiresolution analysis. The developed method can improve the discrimination of functional states collected in longitudinal fMRI studies.

## Introduction

The global brain dynamics generated at rest consists of many superimposed patterns of activity known as resting-state networks (RSNs). RSNs are reproducible in the large samples of healthy subjects^1,2^and are significantly altered in neurological and psychiatric disorders^3,4^. The large-sample assessment of brain resting-state (RS) activity showed more robust results than task-induced fMRI, implying that RS functional connections could serve as biomarkers of brain disorders^5,6^. Thus, RS activity is crucial for understanding the brain activity’s dynamics and organization in health and disease.

Though RSNs are defined based on the absence of goal-directed behavior or salient stimuli, an important question is whether their activity reflects possible residual changes in rest after preceding task-induced activity. This question is especially relevant for understanding the mechanisms of anxiety and post-traumatic stress disorders (PTSD)^7^. Indeed, several fMRI studies showed increased functional connectivity (FC) of some areas in RS after fear conditioning^8–12^. RS fMRI data indicate a high correlation between anxiety and changes in FC^13,14^, as well as an increased FC in patients with PTSD between brain regions that are key for processing emotional stimuli^15^, even years after the stress exposure^16^.

However, the high dimensionality of the fMRI data complicates their conventional analysis based on statistical parametric mapping^17^of changes in the level of blood oxygenation dependent signal (BOLD)^18^. The high dimensionality is especially problematic in the absence of an initial hypothesis that reduces dimensionality by focusing only on regions of interest.

Moreover, in conditions of the high dimensionality of data, valuable results may be lost due to the correction for multiple comparisons in the absence of pre-selected regions of interest. The loss of potential markers is especially crucial when comparing individual longitudinal data or comparing small samples of data in rare clinical diseases. Besides, personalized approaches come to the fore in clinical neuroscience, as it becomes essential to reveal the differences in the activity of the individual’s brain rather than the statistical difference between groups to clarify the diagnosis, the effectiveness of treatment, and rehabilitation. From this point of view, data dimensionality reduction techniques could improve the efficiency of discriminating and classifying brain activity changes at both the individual and group levels, especially in the case of a data-driven approach.

It is believed that almost any high-dimensional data can be effectively nested in the space of significantly lower dimensionality - the so-called intrinsic manifold. When searching for such an internal space, it is usually assumed that there is no a priori data on the positions of points on the intrinsic manifold, that is, the data are unlabeled. The Laplacian eigenmaps (LE) method^19^is a tool developed to solve this problem. It has been used to cluster and classify neuronal activity^20,21^and applied to fMRI data to highlight areas of interest^22^, improve the accuracy of functional network discrimination^23^, analyze fMRI data using diffusion-based spatial priors^24^ and classify brain activity in Alzheimer’s disease^25^. A thorough investigation of machine learning methods in application to resting-state fMRI data can be found in the review of Khosla et al.^26^

In this study, we tested the LE dimensionality reduction method compared to the linear reduction approach via PCA for revealing stimulus-dependent changes in resting-state brain activity before and immediately after fear conditioning and fear extinction in healthy volunteers^27^.

## Methods

### Participants and procedure

We analyzed resting-state fMRI data of 23 healthy right-handed volunteers (23.90 ± 3.93 years old, 8 females) who participated in the study of longitudinal changes in the brain RS functional connectivity such as RS after fear learning and fear memory extinction (RS_FE) in comparison with the initial RS data (RS_0). These data were already used in the previous projects^28^.

### Ethical statement

The study’s protocol followed the Helsinki Declaration’s requirements, and the Ethics Committee of the Institute of Higher Nervous Activity and Neurophysiology of the Russian Academy of Science approved the study. All participants provided written, informed consent before the study.

### Procedure description

The study procedure was as follows: 1) initial resting-state (RS_0) scanning; 2) procedure of fear learning (FL) out of scanner (Pavlovian fear conditioning); 3) fear extinction (FE), and 4) resting-state (RS_FE) scanning after fear extinction. The time between the RS_0 and RS_FE scans was approximately 45 min and between the FE and RS_FE, 1‒2 min. During the RS scanning, participants were asked to remain calm with their eyes closed and to try not to think purposefully. A full description of the experimental procedure can be found in our previous paper^28^. In the current study, we concentrated on analyzing RS1 and RS2 interleaved by FL and FE to separate two RS scan data acquired before and after exposure to emotionally meaningful stimuli.

### Fear learning and fear extinction

The procedure of FL was conducted in a separate room in the behavioral laboratory. The FL consisted of two pseudo-random sequences with a short break between them. For FL, we used a delayed fear-conditioning paradigm with partial negative reinforcement. Three visual (geometric figures) conditioned stimuli were presented. One type of stimuli was always neutral. The presentation of each conditioned stimulus was followed by a white screen for a random duration of 8‒ 12 s with a jitter of 2 s. The other 2 stimuli had the reinforcement probabilities of 70% and 30%, respectively. The unconditional stimulus was mild electrical current stimulation for 500 ms, which was presented immediately after the figure when a white screen appeared. The strength of the stimulation was selected individually to be a tolerable but painful stimulus. Before each stimulus, participants saw a fixation cross lasting 2 s. The duration of each conditional stimulus varied randomly from 4‒8 s with a jitter of 2 s.

The second sequence was the same as the first, except that the probabilities of reinforcement for the conditioned stimuli were changed by 30% and 70%, respectively. The total duration of each FL block was 8 min 54 s.

During the FE session, the same stimuli were presented, but in a different pseudo-random order and with a more-extended overall sequence (10 min) and without US. During FE session, volunteers were asked to expect the unconditional stimulus, but with a different reinforcement rule than in the previous two sessions.

### fMRI data acquisition

The MRI data were collected from the National Research Center Kurchatov Institute (Moscow, Russia) using a 3T scanner (Magnetom Verio, Siemens, Germany) equipped with a 32-channel head coil. The anatomical images were collected using a T1-MPRAGE sequence: TR 1470 ms, TE 1.76 ms, FA 9°; 176 slices with a slice thickness of 1 mm, a slice gap of 0.5 mm, and a 320-mm field of view (FoV) with a matrix size of 320 × 320. Functional images (300 volumes) were collected using a T2*-weighted echo-planar imaging (EPI) sequence having a GRAPPA acceleration factor equal to 4 and the following sequence parameters: TR 2000 ms, TE 20 ms, FA 90°; 42 slices acquired in interleaved order and having a slice thickness of 2 mm, a slice gap of 0.6 mm, and a 200-mm FoV with an acquisition matrix of 98 × 98. Besides, to reduce the EPI’s spatial distortion, the magnitude and phase images were acquired using a field-map algorithm that had the following parameters: TR 468 ms, TE1 4.92 ms, TE2 7.38 ms, an FA 60°, 42 slices, and a 200-mm FOV. The scanner parameters for the RS_0, and RS_FE were the same, with equal session durations of 10 min.

### fMRI data preprocessing

The data of both RS sessions were processed using MELODIC, a part of FSL (FMRIB’s Software Library). We applied the following preprocessing steps: motion correction (MCFLIRT), slice-timing correction using Fourier-space time-series phase-shifting, non-brain removal (BET), spatial smoothing using a Gaussian kernel of FWHM 5 mm, grand-mean intensity normalization of the entire 4D dataset using a single multiplicative factor, high-pass temporal filtering with a removing of the linear trends (Gaussian-weighted, least-squares, straight-line fitting, with sigma = 50.0 s, which equals a cutoff of 0.01 Hz)^29^. The B0-distortion was removed during the inserted B0-unwrapping algorithm. Registration of functional images to the individual anatomical and standard space MNI152 mm3 was conducted using FLIRT^29^. Independent component analysis (ICA) was performed using probabilistic ICA as implemented in MELODIC (v 3.14). For each participant fMRI signal, 38 independent components were extracted. Next, additional denoising of the data was conducted using FIX v1.068 FMRIB’s ICA-based Xnoiseifier^30^ and ICA-based automatic removal of motion artifacts (ICA-AROMA)^31^. First, the AROMA was applied to 15 datasets (five randomly chosen from each RS_0 and RS_FE) in the classification regime to detect motion-related components. The results were then visually inspected to detect additional artifact components, including cerebrospinal fluid (CSF) pulsation in the ventricles. Second, FIX was trained based on the preliminarily classified 15 datasets, and new automatic classification was applied to the remaining 54 sets (23 participants 3 scanning sessions) to detect noisy components, which were filtered out using FIX cleanup mode with the option to clean up the motion confounds (24 regressors). There were no significant within-subject differences in the number of removed components between sessions (mean 19.4 5.06 ICs). Finally, the cleaned data were subjected to filtering to resting-state frequencies of 0.01–0.1 Hz using the “3dTproject” AFNI algorithm.

### Brain parcellation for obtaining BOLD signal

We extracted BOLD signal time series averaged from 245 parcells of the functional resting-state Brainnetom (BN) Atlas^32^. We excluded the BN region 94 from the analysis as this region was absent in some persons who had brain sizes larger than the used field of view. Each mask was converted to individual subject space using FLIRT FSL.

### Analysis framework

Our main approach was to reduce the dimensionality of the data from the number of parcels to an effective one. The value of the BOLD signal indirectly depends on neurons’ activity in the brain structure of interest (Region of Interest - ROI)^18^. If we consider the BOLD response in a single ROI as a separate variable, the brain state at each moment can be represented as an N-dimensional activity vector, where N corresponds to the number of investigated ROIs and depends on the brain parcellation scheme. Along each axis in such space, the value of the signal in a particular area is stored. Thus, a change in the BOLD-signal of the whole brain can be represented as a movement of a point in this multidimensional space.

In our study, 300 BOLD signal values of 245 ROIs (BN parcels) in increments of 2 s corresponded to 300 functional brain images obtained with TR=2s, for each scanning session. In other words, there were 300 points in the high-dimensional space in each session. The task was to separate the brain states related to different sessions (RS_0 and RS_FE) as precisely as possible. For this purpose, 300 points from each session were mixed into a single dataset of 600 points, and class labels were erased. Then the points were divided into two groups employing dimensionality reduction.

### Dimensionality reduction

The LE method^19^ was used to reduce the data dimensionality.

The procedure consisted of two stages:

1. Building a similarity graph based on data in the original high dimensional space.
2. Embedding this graph in a low-dimensional space using its Laplacian eigenvectors.

The first stage was carried out by building the k nearest neighbors graph. For each point, exactly k nearest neighbors (in terms of the Euclidean distance) were calculated if they were connected with the given point. The resulting neighborhood matrix was symmetric. The k parameter was chosen as the minimum possible one to ensure the connectivity of the resulting graph. Such selection preserved the local linearity assumption, which gave sense to the “neighborhood relation” in the initial space.

The second stage was based on the fact that the eigenvectors of the normalized graph Laplacian are also the optimal coordinates for embedding it in a low-dimensional space^19^. To embed N points in the space of dimensionality D, one should take D+1 Laplacian eigenvectors (each of them has length N equal to the number of nodes in the graph). The first eigenvector (the so-called principal eigenvector) was not taken into account in the calculations because its components were constant. In case there were several connected components in a graph, the principal eigenvector had the multiplicity equal to the number of separate graph components. However, such cases were excluded due to the condition of graph connectivity from Stage 1.

To estimate the number of axes of the final space, we used a method of determining the internal dimensionality of data based on the distribution of geodesic distances in a similarity graph^33^. The analysis showed that the number of independent coordinates required to describe the data varies from 5 to 6 for different people (Fig 1).

**Fig. 1.**
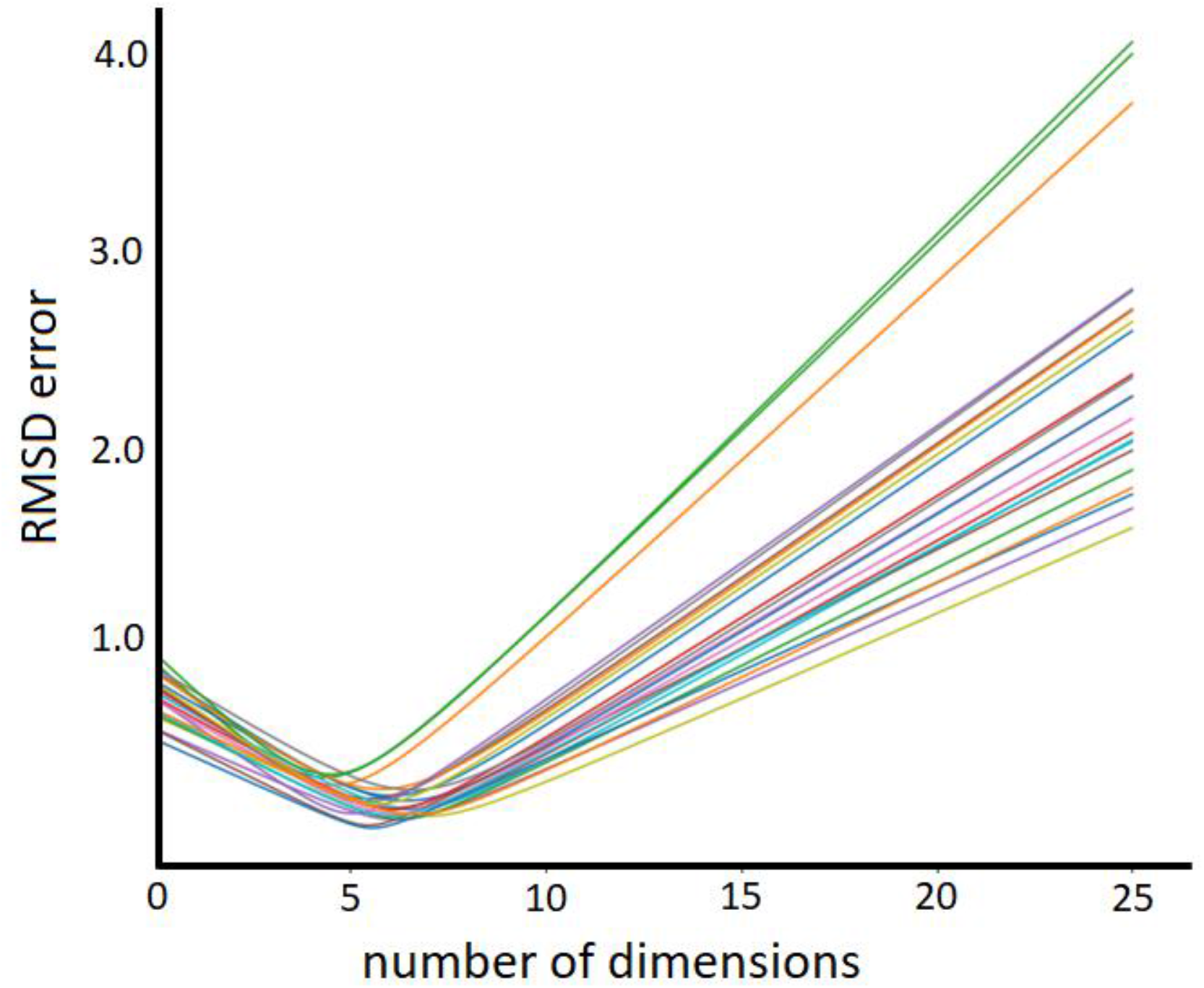
Root mean square deviation between real and theoretic graph geodesic lengths distribution as a function of internal dimensionality tested

Dimensionality reduction was carried out in 2 stages. First, we embedded the data in a 6-dimensional space coinciding with the average internal dimensionality. The resulting embedding points were then used again to construct the similarity graph and project them on a 1-dimensional line using the same algorithm. The same considerations dictated the choice of parameters as at the first reduction of dimensionality. We have chosen k_1_=5, k_2_=20 for KNN graph construction at the first and second stages, respectively.

### Multiscale wavelet transform for smoothing the BOLD signal fluctuations

BOLD-signal has individual variability and non-stationarity. To better distinguish the useful information contained in it, we applied a discrete multiscale wavelet transform^34^. This method has several advantages over the standard Fourier analysis since wavelet base functions are compactly supported and deal well with non-stationary signals.

Following the published recommendations^35^, we selected a wavelet with a sufficiently large number of vanishing moments for the analysis (in our case, it was 10). We chose frequently used Daubechies wavelets as well as the so-called “Symlets” for the analysis.

Multiscale representation provides a hierarchical structure for signal analysis at different “resolutions” and is a natural way of decomposing a signal into several components carrying different aspects of its information content. Such time series representation effectively analyzes the signal’s information content with different levels of detail^36^. At each stage of the algorithm, wavelets with a consistently finer representation of the signal content are generated. Each level of detail represents the signal at a specific resolution.

The first component is a “coarse” version of the signal, which is, in fact, its low-resolution approximation. The other ones are “detail” components that carry information about signal behavior on finer and finer scales, depending on the multiresolution analysis’s depth^36^. A variance of detail components on i-th resolution levels estimates the amount of energy concentrated around the i-th scaling frequency.

During the denoising process, the wavelet coefficients that carry information about the “details” of the signal are filtered at each decomposition stage. There are numerous algorithms for selecting such filtration threshold, adapted to different tasks^37^. After that, filtered detail coefficients are composed again to create a denoised version of an initial signal. The whole process is called “wavelet denoising”.

In this work, a relatively simple and universal VisuShrink method was used^38^. It assumes that the noise in the signal is white and is distributed evenly on all characteristic scales. Under such assumptions, it is reasonable to remove all wavelet coefficients smaller than the noise’s expected maximum amplitude. This amplitude, in turn, can be estimated from the standard deviation of noise and sample size.

Since the form of the “true signal” (and therefore, the noise) is unknown, our estimate of the standard deviation of noise was based on calculating the median of detailed wavelet coefficients^39^.

The time series used in the numerical experiments were two artificially stacked sequences of 300 points each. Therefore, wavelet denoising was performed separately for the first and second half of the combined signal (Fig 2). Shuffling of N-dimensional activity vectors (N stands for the number of ROIs) was not performed as timestamps were not preserved during the construction of a similarity graph. Therefore, the ordering of activity vectors does not affect the final low-dimensional embedding.

**Fig. 2.**
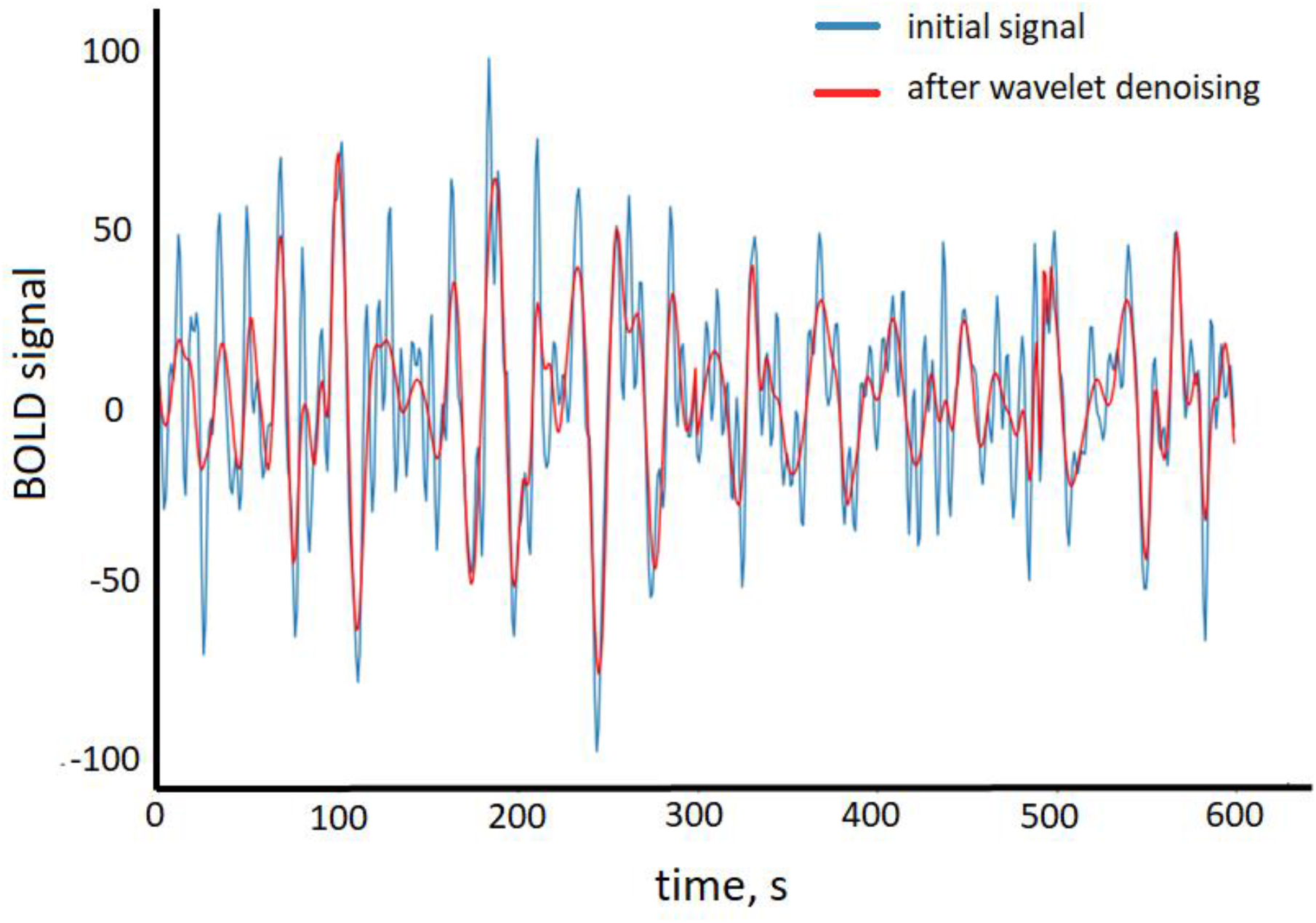
An example of an initial (blue) and filtered (red) BOLD-signal

### Discriminating two resting-state sessions

The ratio of the average distance between points of different classes to the average distance between points of the same class (interclass-intraclass distance, IID) was used as a measure of separation of two classes of points (corresponding to sessions before and after exposure). In this study, we decided to focus on IID analysis because it measures the distance between two populations “in general,” which is of interest.

The standard binary classification quality metrics show how well the algorithm handles test data labeling, provided that the class labels for training data are known. We were also interested in another question - how well the data belonging to different scanning sessions were separated in the new low-dimensional space. This is why IID was chosen as the primary metric.

In addition to IID, the accuracy and ROC-AUC score of the K-means clustering algorithm with k=10 was calculated. The results were cross-validated using ten folds (10 non-overlapping train-test data splits, test set contains 10% of data in each split).

To show the non-linear nature of the stated problem, we compare the LE dimensionality reduction results with PCA ones. We applied PCA on a data matrix of shape 245×600, transforming it to shape TDx600, where TD is target dimensionality (number of principal components taken).

All results were averaged over subjects after removing outliers that stood out from the average by more than 2.5 standard deviations.

## Results

### Comparison of separation accuracy of fMRI data

IID between two sessions after applying PCA was only slightly higher from random: 1.014 ± 0.011 for simple PCA and 1.022 ± 0.015 for PCA with preliminary wavelet denoising. However, wavelet denoising increased accuracy and ROC_AUC score (see fig. S1).

After the dimensionality reduction with the non-linear LE algorithm, it was possible to separate the data points belonging to different scanning sessions. For most subjects (19 out of 23), the low-dimensional projection points split up into two clusters corresponding to the sessions before and after fear learning (Fig.3 A and C). The remaining four subjects did not show such discrimination (Fig.3 B and D), but for all subjects, the IID was significantly higher than a random one (if we assign labels to data points randomly, the IID will not exceed 1.01 ± 0.01). The IID values for individual subjects can be found in the table S1.

**Fig. 3.**
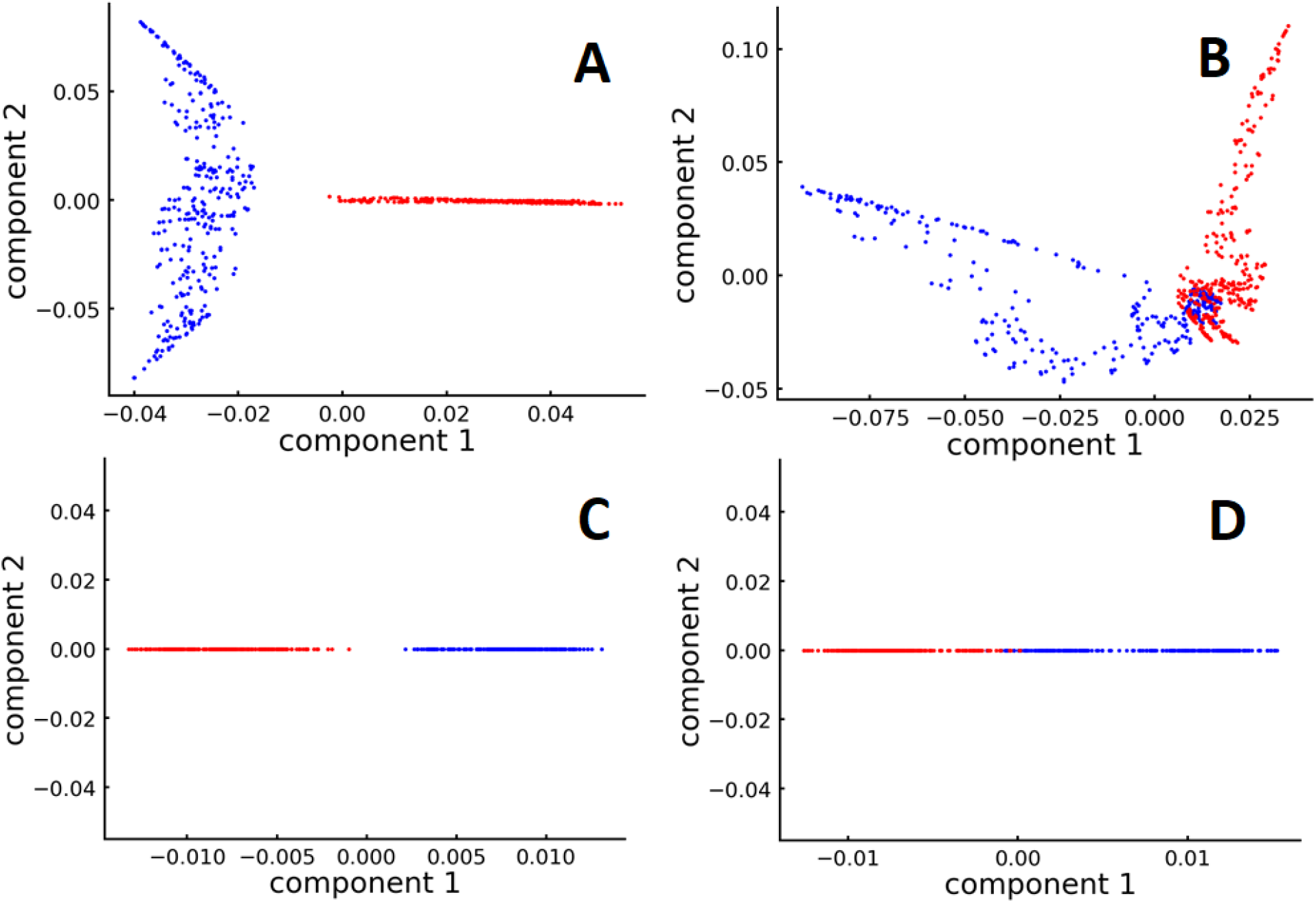
An example of a “good” (left) and “bad” (right) dimensionality reduction result. Red and blue points correspond to data from RS_0 and RS_FE sessions respectively. A, B: two-dimensional projections of an intermediate 6-dimensional embedding. C, D: projections on a one-dimensional line from an intermediate embedding. IID values: 8.23 for fig. 3C, 2.26 for fig. 3D

### Wavelet denoising improves the separation of BOLD signal data from different sessions

After applying a multiscale wavelet transform, shrinking the detail coefficients, and collecting the truncated components back into a time series, we obtained denoised versions of our BOLD signals (see Methods). After applying dimensionality reduction to this new filtered data, IID increased for the majority of subjects, see Fig. 4 (average increase of 52% ± 30% for a single dimensionality reduction (dim 245 -> dim 1, see Fig. 4) and 250% ± 90% for a two-step dimensionality reduction (dim 245 -> dim 6 -> dim 1, see Fig. 6). Detailed results can be found in the table S1.

**Fig. 4.**
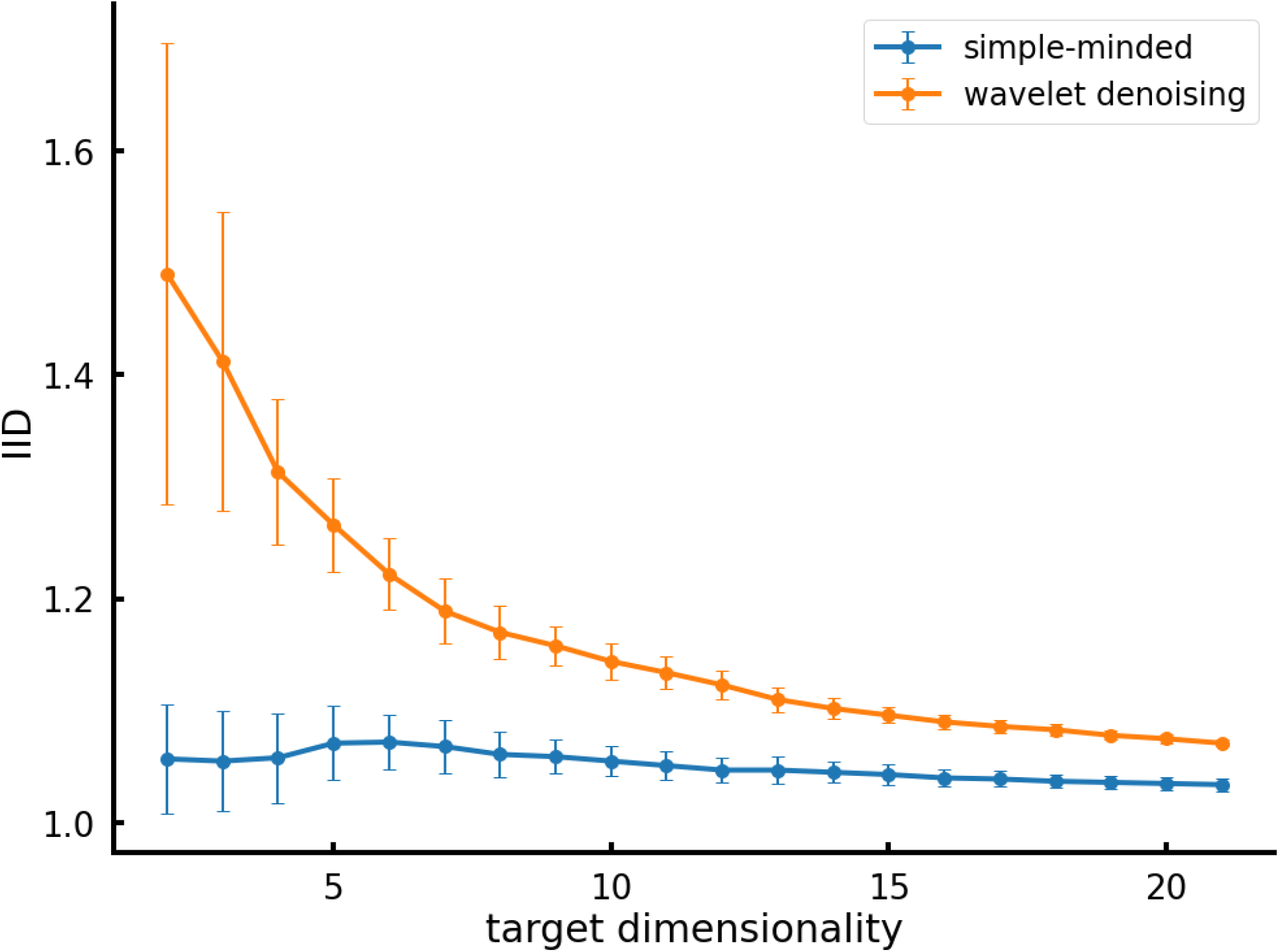
Improved separation between two RS scan sessions after applying wavelet denoising. Dimensionality was reduced in one step (dim 245 -> target dim) with LE method. Bars represent standard deviations.

Using BOLD preprocessing with wavelets allowed us to observe two clear clusters corresponding to functional states before and after exposure for many subjects (Figs. 3 A and C). Later on, in all further analysis, we used wavelet denoising for the BOLD signal.

Experimenting with different dimensions of the final space, we concluded that a better separation (IID = 1.82±0.5) was achieved when projecting to a line (1D space), see Fig. 5. At the same time, applying PCA dimensionality reduction resulted in much lower IID≈1,1. The same results apply to classification accuracy and ROC_AUC score (see fig. S2).

**Fig. 5.**
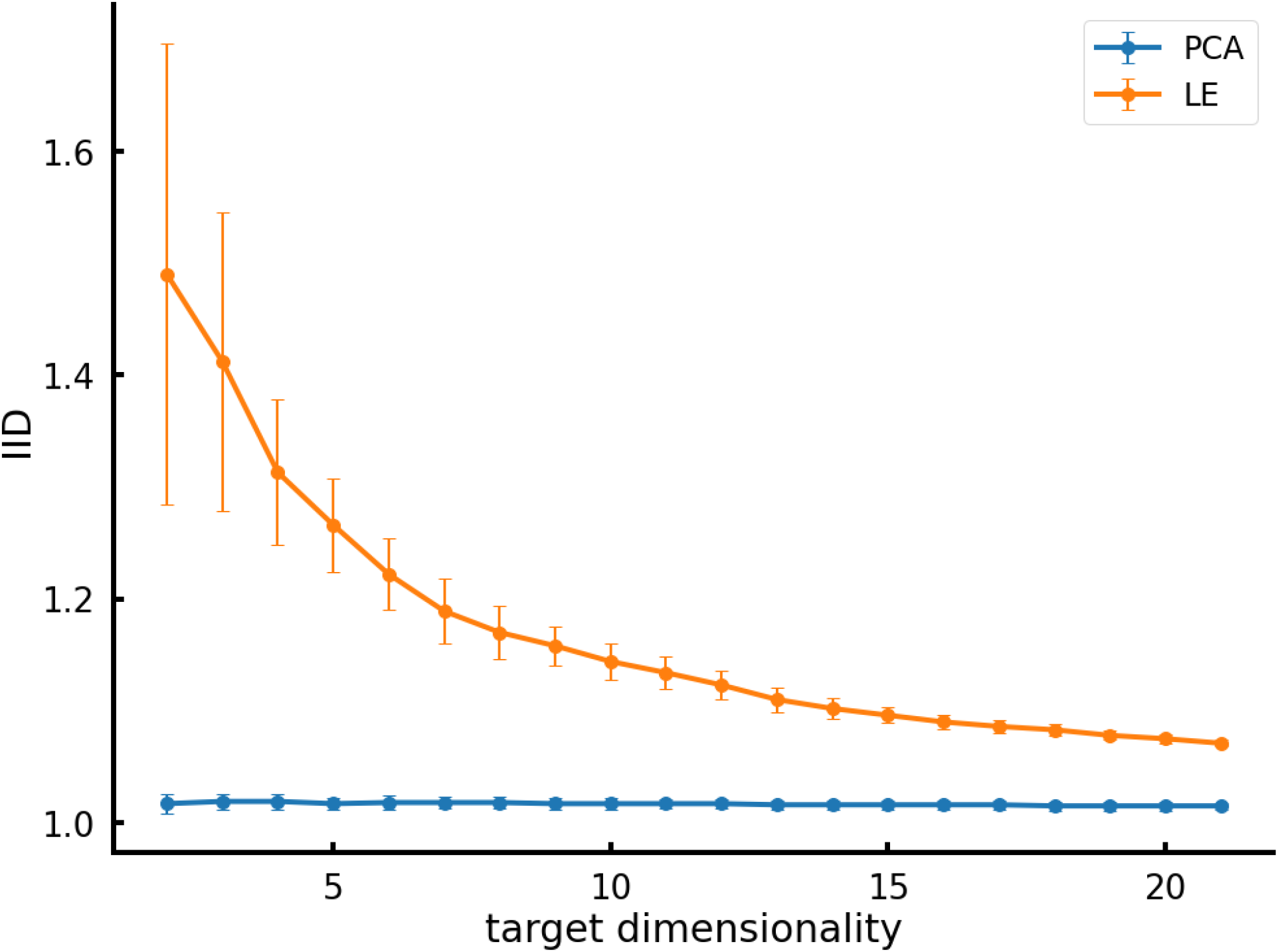
Dependence of the average IID on the dimensionality of the final space to which the embedding was performed (dim 245 -> target dim). The blue line represents PCA results; the yellow line stands for LE.

The average IID increased by ≈100% after we started to use a preliminary reduction of data dimensionality to the internal one (6.1 ± 0.6), see Fig 6. This effect seems to be due to the peculiarity of LE method, which tries to preserve proximity in a low-dimensional space only for the points connected by edges in the similarity graph. Preliminary reduction of dimensionality allows an additional separation of two groups of points belonging to different scanning sessions.

**Fig. 6.**
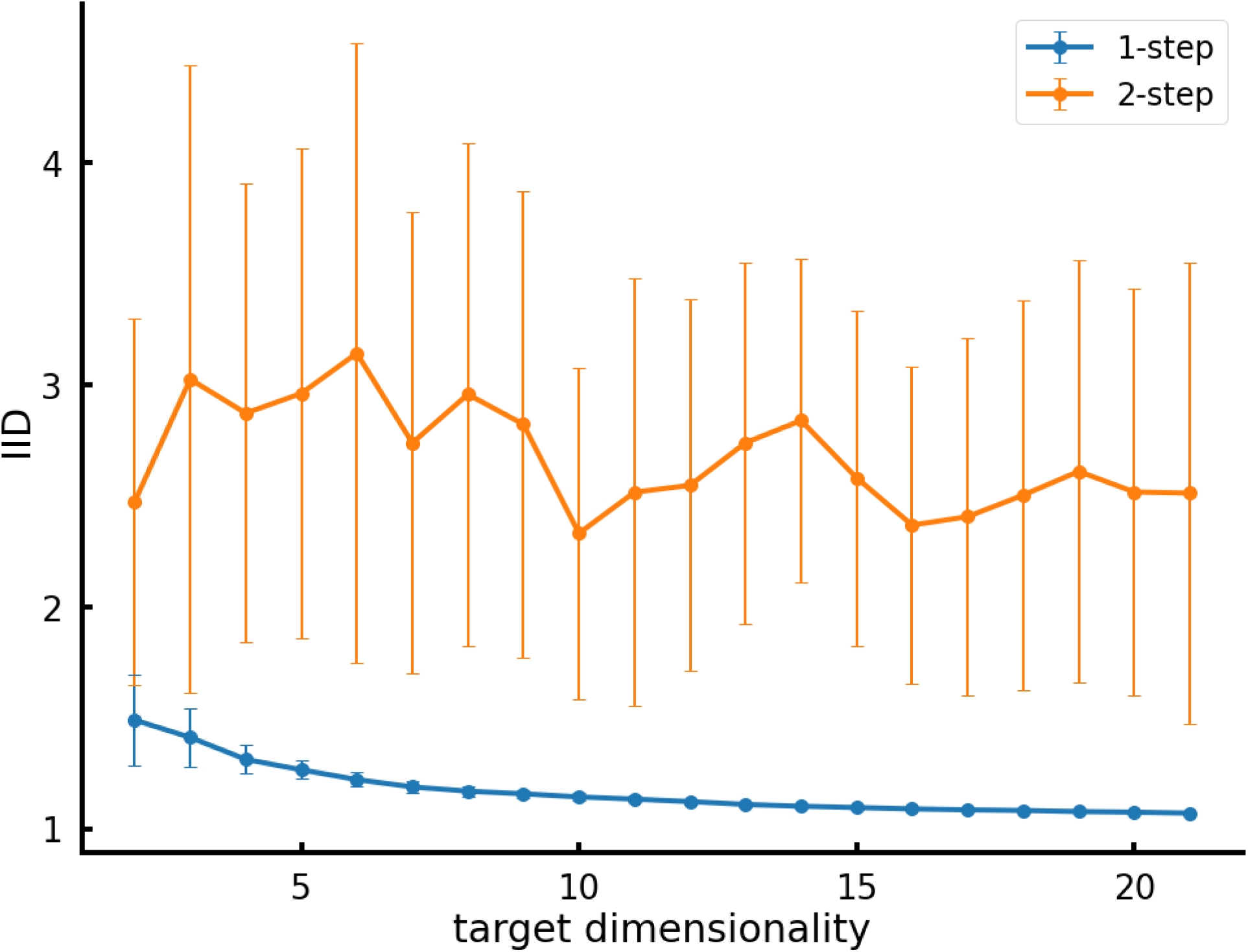
Comparison of IIDs for direct LE dimensionality reduction (245->TD, blue line) and 2 steps dimensionality reduction (245 -> target dim -> 1, yellow line) depending on the target dimensionality.

## Discussion

Our study aimed to develop a dimensionality reduction method to classify individual differences in BOLD-signal at rest before and after salient stimulus exposure. BOLD-signal values used from the 245 ROIs are not fully independent, and the number of axes required for sufficient data description is significantly lower. However, the interdependencies between the different components of a multidimensional signal can be complex and non-linear. Therefore, we decided to use dimensionality manifold learning methods to identify the data structure in their intrinsic manifold.

For the first time, we show that it is possible to distinguish brain activity at rest before and after stimulus exposure using fMRI data at the level of individual subjects with the high precision.

The PCA linear dimensionality reduction method was not successful in this task. Low-dimensional representations returned by linear methods are associated with input vectors of linear transformation features^19^. If the data belong to some linear subspace of the original high-dimensional representation (or are located close to it), the linear methods may create a good embedding into a low-dimensional space. However, they will fail if the data lie on a non-linear manifold because the surface geometry becomes much more complex. Therefore, the LE method was used in this work to restore the internal geometry of data.

When choosing a dimensionality reduction algorithm, we were guided first of all by its ability to restore the “internal manifold” of our data. Among non-linear methods LE was chosen because of its small computational complexity and the clear geometric meaning of the reduced space coordinates - they ability to dictate the optimal division of the graph into non-intersecting clusters^40^.

Our task was not just to develop a classifier that would distinguish data from the first and the second sessions. We wanted to build a low-dimensional embedding of the data that would naturally separate the two data groups. Therefore we focused on the manifold learning algorithms. This approach was successfully adopted in recent papers: the local linear embedding algorithm served as a useful feature extraction procedure that helped to recognize psychiatric disorders^41^. Manifold discovery using LE was performed on fMRI images^22^. Being applied to fMRI images, this technique facilitated the discrimination of structures of interest. Manifold learning methods have been applied to functional brain networks^42^ to construct optimal low-dimensional embeddings.

BOLD signal preprocessing through wavelet denoising helped to improve the separation of two data groups significantly. The best wavelet we tested was Daubechies 10. Most likely, this is due to its resemblance to the characteristic form of a BOLD signal, as well as to the high number of vanishing moments^35^.

The optimal number of levels (n) for multiresolution analysis was 3: this seems to be because with n<=2 the BOLD-signal was not filtered on a large scale. The frequency range often interpreted in resting fMRI is 0.008-0.15Hz^35^. In our case, we used a band-pass filter of 0.01-0.1 Hz. Scanning was performed with TR = 2s; thus, scale 2 coefficients correspond to the range 0.025-0.05 Hz, and scale 3 coefficients correspond to the range 0.0125-0.025 Hz. It seems that the wavelet coefficients in these two frequency bands are essential for separating brain states before and after exposure. A detailed comparison of the results for different wavelets for individual subjects can be found in the table S1.

An important issue when reducing the dimensionality of data is the optimal number of axes of the embedding. In this study, we found that for all subjects, the internal dimensionality of the original 245-dimensional data ranges from 5 to 6. This finding is most likely associated with the high degree of correlation between BOLD signals in different brain regions. In the future, analysis of these several non-linear components may shed light on the integration processes in the brain and the architecture of its functional networks.

The best separation of data from two sessions was obtained in a 1D projection. In this case, the only coordinate specifying the position of a point on the line is the component of the first nontrivial Laplacian eigenvector (the so-called Fiedler vector), which is responsible for cutting the graph into two optimal halves^40^.

However, this does not mean that the data can be well described with a single variable. Instead, we have revealed the meaning of only one, albeit the most important, coordinate axis in the latent space. The meaning of the other variables remains to be recognized.

As a limitation of the results, we should mention that, one of the main problems of manifold learning methods is the interpretation of latent space axes, which is often done empirically, without any initial hypothesis. Hence, the work on “mapping” the space of hidden variables^43^ is critical and can provide much new information about the data and its hidden variables. Our findings are consistent with the previous data^44–46^, the main difference being a successful application of the LE dimensionality reduction to separate the brain RS activity before and after meaningful stimulus exposure.

## Conclusions

Our study demonstrated that the Laplacian eigenmaps manifold learning method applied to fMRI data could reveal differences between the RS functional brain activity before and after exposure of subject to a salient stimulus. Our findings also demonstrate that RS functional brain networks retain effects of stimuli exposure even after a time lag.

## Author Contributions

Those who conceived and designed the study include NP, AT, OM and KA. AT performed the experiments. AT and NP analyzed the data. NP and OM wrote the paper.

## Acknowledgements

We are grateful to S. Nechaev and A. Gorsky for the important comments.

## References

1. van den Heuvel MP, Hulshoff Pol HE. Exploring the brain network: A review on resting-state fMRI functional connectivity. Eur Neuropsychopharmacol. 2010. doi:10.1016/j.euroneuro.2010.03.008

2. Martynova O V., Sushinskaya-Tetereva AO, Balaev V V., Ivanitskii AM. Correlation between the Functional Connectivity of Brain Areas Active in the Resting State with Behavioral and Psychological Indicators. Neurosci Behav Physiol. 2017. doi:10.1007/s11055-017-0520-1

3. Hohenfeld C, Werner CJ, Reetz K. Resting-state connectivity in neurodegenerative disorders: Is there potential for an imaging biomarker? NeuroImage Clin. 2018. doi:10.1016/j.nicl.2018.03.013

4. Sheffield JM, Barch DM. Cognition and resting-state functional connectivity in schizophrenia. Neurosci Biobehav Rev. 2016. doi:10.1016/j.neubiorev.2015.12.007

5. Satterthwaite TD, Baker JT. How can studies of resting-state functional connectivity help us understand psychosis as a disorder of brain development? Curr Opin Neurobiol. 2015. doi:10.1016/j.conb.2014.10.005

6. Barch DM. Resting-state functional connectivity in the human connectome project: Current status and relevance to understanding psychopathology. Harv Rev Psychiatry. 2017. doi:10.1097/HRP.0000000000000166

7. Graham BM, Milad MR. The study of fear extinction: Implications for anxiety disorders. Am J Psychiatry. 2011. doi:10.1176/appi.ajp.2011.11040557

8. Hahn A, Stein P, Windischberger C, et al. Reduced resting-state functional connectivity between amygdala and orbitofrontal cortex in social anxiety disorder. Neuroimage. 2011. doi:10.1016/j.neuroimage.2011.02.064

9. Prater KE, Hosanagar A, Klumpp H, Angstadt M, Phan KL. Aberrant amygdala-frontal cortex connectivity during perception of fearful faces and at rest in generalized social anxiety disorder. Depress Anxiety. 2013. doi:10.1002/da.22014

10. Baeken C, Marinazzo D, Van Schuerbeek P, et al. Left and right amygdala - Mediofrontal cortical functional connectivity is differentially modulated by harm avoidance. PLoS One. 2014. doi:10.1371/journal.pone.0095740

11. Rus OG, Reess TJ, Wagner G, Zimmer C, Zaudig M, Koch K. Functional and structural connectivity of the amygdala in obsessive-compulsive disorder. NeuroImage Clin. 2017. doi:10.1016/j.nicl.2016.12.007

12. Jung YH, Shin JE, Lee YI, Jang JH, Jo HJ, Choi SH. Altered amygdala resting-state functional connectivity and hemispheric asymmetry in patients with social anxiety disorder. Front Psychiatry. 2018. doi:10.3389/fpsyt.2018.00164

13. Kim MJ, Gee DG, Loucks RA, Davis FC, Whalen PJ. Anxiety Dissociates dorsal and ventral medial prefrontal cortex functional connectivity with the amygdala at rest. Cereb Cortex. 2011. doi:10.1093/cercor/bhq237

14. Belleau EL, Pedersen WS, Miskovich TA, Helmstetter FJ, Larson CL. Cortico-limbic connectivity changes following fear extinction and relationships with trait anxiety. Soc Cogn Affect Neurosci. 2018. doi:10.1093/scan/nsy073

15. Zhou Y, Wang Z, Qin L di, et al. Early Altered Resting-State Functional Connectivity Predicts the Severity of Post-Traumatic Stress Disorder Symptoms in Acutely Traumatized Subjects. PLoS One. 2012. doi:10.1371/journal.pone.0046833

16. Brown VM, Labar KS, Haswell CC, et al. Altered resting-state functional connectivity of basolateral and centromedial amygdala complexes in posttraumatic stress disorder. Neuropsychopharmacology. 2014. doi:10.1038/npp.2013.197

17. Fristen KJ, Frith CD, Fletcher P, Liddle PF, Frackowiak RSJ. Functional topography: Multidimensional scaling and functional connectivity in the brain. Cereb Cortex. 1996. doi:10.1093/cercor/6.2.156

18. Poldrack RA, Fletcher PC, Henson RN, Worsley KJ, Brett M, Nichols TE. Guidelines for reporting an fMRI study. Neuroimage. 2008. doi:10.1016/j.neuroimage.2007.11.048

19. Belkin M, Niyogi P. Laplacian eigenmaps for dimensionality reduction and data representation. Neural Comput. 2003. doi:10.1162/089976603321780317

20. Sun G, Zhang S, Zhang Y, et al. Effective dimensionality reduction for visualizing neural dynamics by laplacian eigenmaps. Neural Comput. 2019. doi:10.1162/neco_a_01203

21. Rubin A, Sheintuch L, Brande-Eilat N, et al. Revealing neural correlates of behavior without behavioral measurements. Nat Commun. 2019. doi:10.1038/s41467-019-12724-2

22. Thirion B, Faugeras O. Nonlinear dimension reduction of fMri data: The laplacian embedding approach. In: 2004 2nd IEEE International Symposium on Biomedical Imaging: Macro to Nano.; 2004. doi:10.1109/isbi.2004.1398552

23. Liu C, JaJa J, Pessoa L. LEICA: Laplacian eigenmaps for group ICA decomposition of fMRI data. Neuroimage. 2018. doi:10.1016/j.neuroimage.2017.12.018

24. Harrison LM, Penny W, Daunizeau J, Friston KJ. Diffusion-based spatial priors for functional magnetic resonance images. Neuroimage. 2008. doi:10.1016/j.neuroimage.2008.02.005

25. Hu C, Sepulcre J, Johnson KA, Fakhri GE, Lu YM, Li Q. Matched signal detection on graphs: Theory and application to brain imaging data classification. Neuroimage. 2016. doi:10.1016/j.neuroimage.2015.10.026

26. Khosla M, Jamison K, Ngo GH, Kuceyeski A, Sabuncu MR. Machine learning in resting-state fMRI analysis. Magn Reson Imaging. 2019. doi:10.1016/j.mri.2019.05.031

27. Martynova O, Tetereva A, Balaev V, Portnova G, Ushakov V, Ivanitsky A. Longitudinal changes of resting-state functional connectivity of amygdala following fear learning and extinction. Int J Psychophysiol. 2020. doi:10.1016/j.ijpsycho.2020.01.002

28. Tetereva A, Kartashov S, Ivanitsky A, Martynova O. Variance and Scale-Free Properties of Resting-State Blood Oxygenation Level-Dependent Signal After Fear Memory Acquisition and Extinction. Front Hum Neurosci. 2020. doi:10.3389/fnhum.2020.509075

29. Jenkinson M, Bannister P, Brady M, Smith S. Improved optimization for the robust and accurate linear registration and motion correction of brain images. Neuroimage. 2002. doi:10.1016/S1053-8119(02)91132-8

30. Griffanti L, Salimi-Khorshidi G, Beckmann CF, et al. ICA-based artefact removal and accelerated fMRI acquisition for improved resting state network imaging. Neuroimage. 2014. doi:10.1016/j.neuroimage.2014.03.034

31. Pruim RHR, Mennes M, van Rooij D, Llera A, Buitelaar JK, Beckmann CF. ICA-AROMA: A robust ICA-based strategy for removing motion artifacts from fMRI data. Neuroimage. 2015. doi:10.1016/j.neuroimage.2015.02.064

32. Fan L, Li H, Zhuo J, et al. The Human Brainnetome Atlas: A New Brain Atlas Based on Connectional Architecture. Cereb Cortex. 2016. doi:10.1093/cercor/bhw157

33. Granata D, Carnevale V. Accurate Estimation of the Intrinsic Dimension Using Graph Distances: Unraveling the Geometric Complexity of Datasets. Sci Rep. 2016. doi:10.1038/srep31377

34. Abry P, Baraniuk R, Flandrin P, Riedi R, Veitch D. Multiscale nature of network traffic. IEEE Signal Process Mag. 2002. doi:10.1109/79.998080

35. Zhang Z, Telesford QK, Giusti C, Lim KO, Bassett DS. Choosing wavelet methods, filters, and lengths for functional brain network construction. PLoS One. 2016. doi:10.1371/journal.pone.0157243

36. Mallat SG. A Theory for Multiresolution Signal Decomposition: The Wavelet Representation. IEEE Trans Pattern Anal Mach Intell. 1989. doi:10.1109/34.192463

37. Luo G, Zhang D. Wavelet Denoising. In: Advances in Wavelet Theory and Their Applications in Engineering, Physics and Technology.; 2012. doi:10.5772/37424

38. Donoho DL. De-Noising by Soft-Thresholding. IEEE Trans Inf Theory. 1995. doi:10.1109/18.382009

39. Mallat S. A Wavelet Tour of Signal Processing.; 2009. doi:10.1016/B978-0-12-374370-1.X0001-8

40. Shi J, Malik J. Normalized cuts and image segmentation. IEEE Trans Pattern Anal Mach Intell. 2000. doi:10.1109/34.868688

41. Sidhu G. Locally Linear Embedding and fMRI Feature Selection in Psychiatric Classification. IEEE J Transl Eng Heal Med. 2019. doi:10.1109/JTEHM.2019.2936348

42. Qiu A, Lee A, Tan M, Chung MK. Manifold learning on brain functional networks in aging. Med Image Anal. 2015. doi:10.1016/j.media.2014.10.006

43. Liu Y, Jun E, Li Q, Heer J. Latent space cartography: Visual analysis of vector space embeddings. Comput Graph Forum. 2019. doi:10.1111/cgf.13672

44. Bahrami M, Lyday RG, Casanova R, Burdette JH, Simpson SL, Laurienti PJ. Using Low-Dimensional Manifolds to Map Relationships Between Dynamic Brain Networks. Front Hum Neurosci. 2019. doi:10.3389/fnhum.2019.00430

45. Hong SJ, Xu T, Nikolaidis A, et al. Toward a connectivity gradient-based framework for reproducible biomarker discovery. Neuroimage. 2020. doi:10.1016/j.neuroimage.2020.117322

46. Billings JCW, Medda A, Shakil S, et al. Instantaneous brain dynamics mapped to a continuous state space. Neuroimage. 2017. doi:10.1016/j.neuroimage.2017.08.042

